# Differentiation in the body color of *Creobroter gemmata* under red and green background at an early stage

**DOI:** 10.1101/2025.03.05.641766

**Authors:** Shao Zhi George Liu, Renzhi Ma

**Affiliations:** New York University

**Keywords:** *Creobroter gemmata*, mantis, background color, camouflage, pigmentation, image analysis, body color

## Abstract

Praying mantises (*Mantodea*) are the master of camouflage. They passively prey on any small animals that intrude their striking distance with mantises’ powerful grasping legs. *Creobroter gemmate* belongs to the family of flower mantis (*Hymenopodidae*), in which most species mimic flowers. Their body color consequently becomes various. Many factors can affect the body color of praying mantises, including moisture and temperature. However, there is little research showing the relation between the background color and their body color. This experiment determined the influence of background color on *Creobroter gemmata*’s body color by raising nymphs in the red and green background and using image analysis. The results indicate that *Creobroter gemmate* will change their body color according to the background, and the differentiation is more significant in the early stages. The discovery shows how background color plays a role in insects’ development and brings forward a mechanism of insects’ camouflage, which is worth further studying.

## 1. Introduction

Mimicry is a usual appearance for many species of arthropods, both predators and prey (Marc & Jérôme, 2009). Entomologists have investigated the masterful mimicry of praying mantis since the 20th century (Edmunds, 1999). Species of *Mantodea* imitate fresh or dead leaves, grasses, branches, or flowers, according to their habitats. Some of these mantis species are able to change their color and pattern of patches within their life cycle in order to adapt to seasonal and environmental changes (Mukherjee, *et al*., 1995). The type of vegetation in the surrounding environment, the intensiveness of sunlight, and the months in a year could all be the factors that differentiate mantis’ body color.

Having diverse body shapes and colors, praying mantis are favored by many scientists and insects lovers. People are willing to manipulate praying mantis’ body color to obtain more beautiful pets’ categories or get a deeper insight into the mechanism of insects’ camouflage. The differentiation of body color is also helpful for taxonomists to determine new species (Svenson, *et al*., 1915). Tracing the body color of a praying mantis in an artificial environment by using digital techniques may provide scientists some clues about the mantis’ coloration mechanism.

In the wild, a well-selected body color that resembles surrounding environments gives individuals more chance to escape from predators, and praying mantis possibly actively prefer microhabitats that match their coloration (Battiston, *et al*., 2010; Hurd & Eisemberg, 1984). Current studies about the body color shift in *Orthopterodea* are mostly focused on *Orthoptera* insects, such as grasshopper and locust (Edelaar, *et al*., 2017; Tanaka, 2001). These studies point out that the change in body color also depends on the endocrine system, mechanical stimulation, and substrate color. Substrate color, which is also known as the colored light reflected from the background environment, was considered as one of the factors of the change in body and pattern color of the *Orthopterodea* insects (Hochkirch, *et al*., 2008). As the order belongs to *Orthopterodea* as well, *Mantodea* is also supposed to be capable of mimicking a diverse environment and has a variety of body colors. However, old research shows that young nymphs of *Mantis religiosa* from the family of *Mantidae* did not increase their green pigment formation in a green background (Okay, 1953).

Although the experiment of *Mantis religiosa* shows a negative result, there are still many features that indicate that some species of praying mantis are likely to tie up their body color with the background color. Praying mantis, especially species in the family *Hymenopodidae*, are experts in mimicking flowers and withered leaves. The orchid mantis *Hymenopus coronatus* has been hypothesized to mimic a flower corolla. They can change their body color from pink to white through growing up by molting (O’Hanlon, *et al*., 2014). This skill is probably out of the seasonal transition of surrounding orchid flowers. As a member of *Hymenopodidae, Creobroter gemmate*, also known as eyespot mantis, also mimics flowers. It possesses high diversity of body color. Wild individuals’ appearance with green, yellow, red even pink were observed (Schwarz, 2015). The species of *Creobroter gemmata* has green and white abdominal patches, which help them build up the shape and color of their disguise. As nymphs, the flowery disguise is more intense than the adults. They usually hang upside down on flora to create an invisible disguise and fool the potential prey of flying insects which eager to suck in a drop of nectar.

This research hypothesizes that the background color of a grow-up environment will differentiate the body color of *Creobroter gemmata*. The implication of this study would lead to a better understanding of the mechanism of insects’ imitation deeper and exploit a new region of insects’ multiple coloration plasticity. To find out how background color determines the gradual change in mantis’ body color, this research analyzed the body color of individuals in *Creobroter gemmata* and compared the pictures of the mantis grown in red, green, and white background from the 1st instar to the 5th.

## 2. Method

### 2.1 Breeding device

In the experiment, all mantis were bred in an incubator with constant temperature, moisture, and photo-period (26 Celsius degree and 60 percent of moisture). Every mantis nymph was fed in a translucent plastic box individually. Sixty individuals of mantis nymphs were raised and separated into three groups. The 60 mantis nymphs were averagely divided into three groups, each with 20 individuals. These three groups correspond to boxes with the red substrate, green substrate, and without substrate. The group without substrate is in the white box, considered as the control group.

The size of the boxes is 6.5cm*6.5cm*4.5cm shown as Fig. 1. A white fibrous paste is set for mantis nymphs to grip and exuviate on each top of the box. All the mantis were fed by fruit fly. Nymphs at 1st instar was fed by *Drosophila melanogaster*, due to its smaller size, and other nymphs are fed by *Drosophila pseudoobscura*. All nymphs were fed once a week, and ensured to be well fed by fruit flies and sufficient pure water (Yager, *et al*., 1999).

**Fig. 1.**
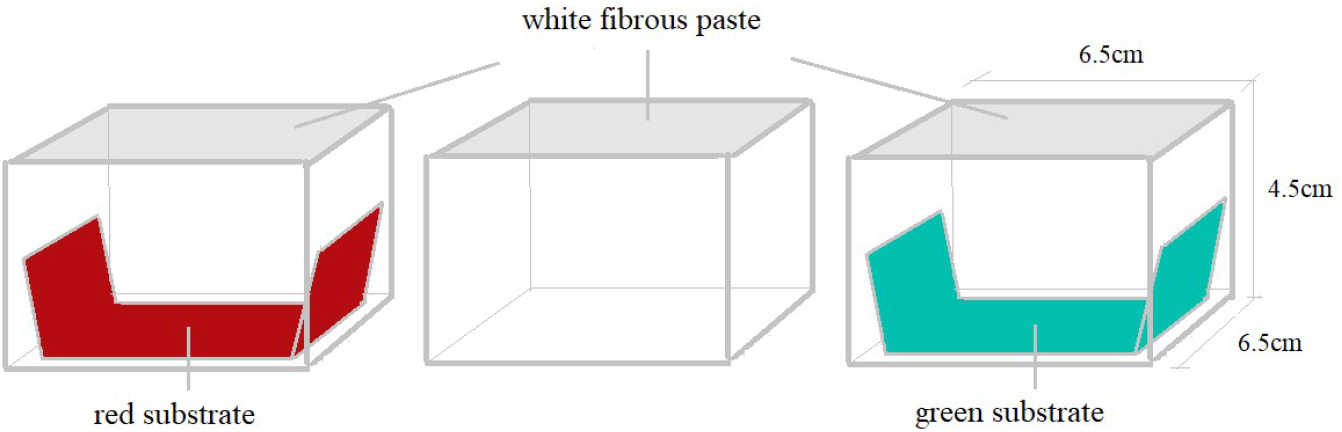
The picture shows the devices used to keep the nymphs. They are transparent plastic boxes (6.5cm*6.5cm*4.5cm) with white fibrous pastes on the top. The red box and the green box contain a trapezoid reflective plastic color substrate at the bottom.

### 2.2 Background color

The substrate background is used to reflect the colored light on the praying mantis. The environment is capable of scattering the colored light on every aspect of the praying mantis so that the red and green plastic plates are inverted to a trapezoid shape, known as the red/green substrate. The RGB value of the red substrate plastic plate is (#B50C11) Red: 181, Green: 12, Blue: 17; the green substrate plastic plate is (#01BFAE) Red: 1, Green: 191, Blue: 174 (Fig. 2). Plastic plates are highly reflective, immersing the entire box in the substrate color. The background colors chosen are red and green because the wild individuals of the *Creobroter gemmata* are mainly red, brown, or green, which is highly conform with the habitat with foliage and flower.

**Fig. 2.**
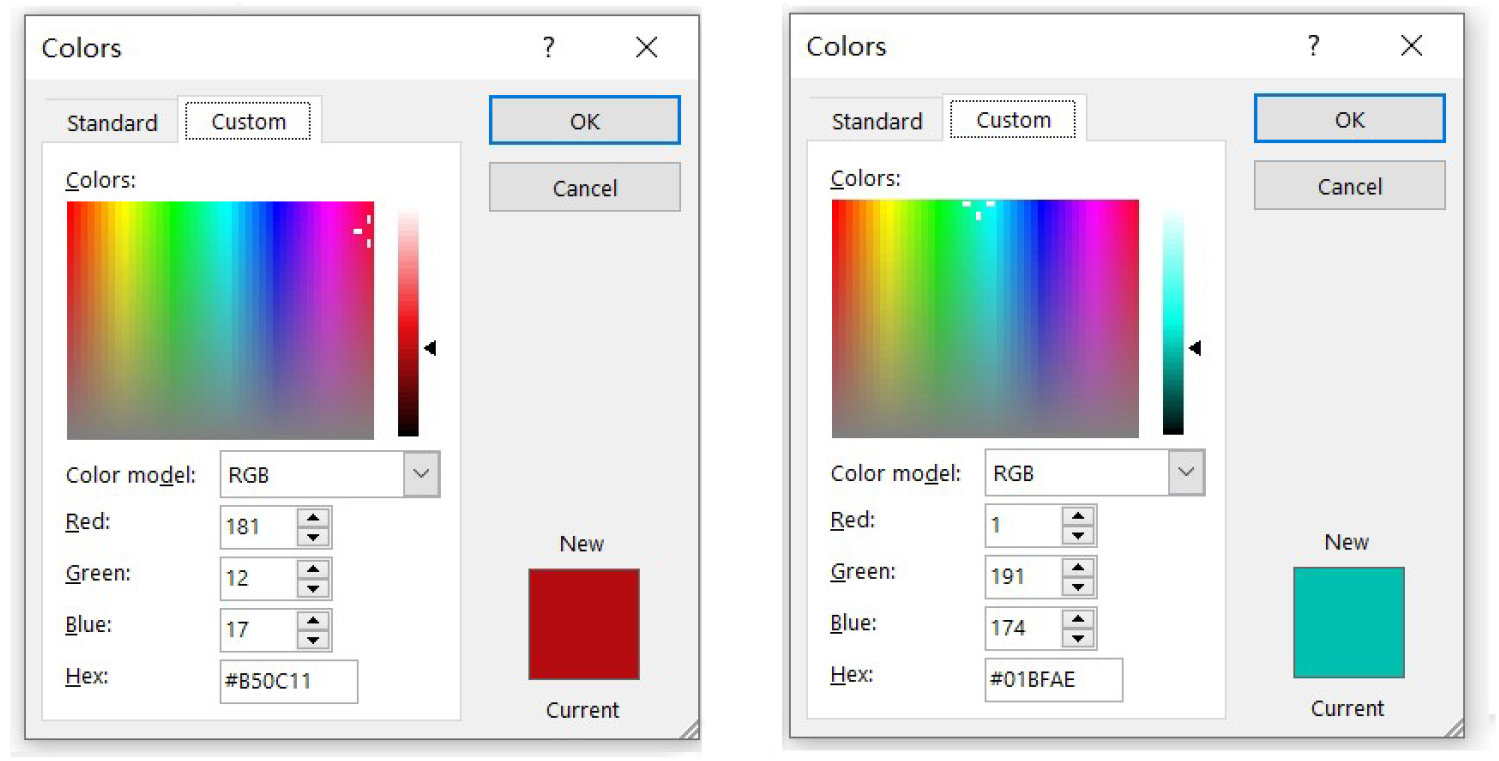
The picture shows the all three RGB values and the appearance of two substrate colors.

### 2.3 The source of mantis nymphs

To observe the whole process of the growth in the colored background, the nymphs have been separated into individual boxes since they were hatched. The younger the nymphs are, the lower the survival rate is. Three oothecae were incubated at different timing to ensure the quantity. The oothecae were hatched in 26 Celcius degrees and about 60 percent of moisture as well.

### 2.4 Image acquisition

To investigate the body color’s gradual transition, individual’s different aspects at each instar are needed to collect. Except for the 1^st^ and the 5^th^ instar, 5 mantis individuals are randomly selected from the total of 20 mantes to take the photo. The 1^st^ and the 5^th^ instar collected all 20 samples from each group. Three representative parts of the mantis body were chosen for taking photos: The grasping leg, the pronotum, and the sterna. Each part has a unique pattern of patches and is evidently bright-colored. All of the photos taken have a magnification of 400% under the microscope, and the lights are illuminated from the same directions. The mantis nymph is placed on a glass plate. By Rotating and adjusting the position of the glass plate, pictures of specific body parts are taken. The position and the body size of mantis nymphs are unpredictable, so the shape photos were not completely identical (Martineau, *et al*., 2017).

### 2.5 Data processing

After collected the images, Data is analyzed by RGB values. A python program will run through every pixel in the image, which will select those with color in an acceptable range of RGB values. Then, it will calculate the average RGB values of all selected pixels.

The pixels which are overly bright or dark will be excluded by the program. Specifically, if all three values of the RGB of a pixel are higher than 230 or lower than 25, the pixel will not be added into account. Thereafter, the values will be calculated by a formula and show an index that indicates the extent of green and red body color. Pictures in the same group will finally be used to reduce a certain value. Then, the indexes used to represent the extent of red and green are calculated for each group:

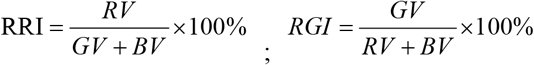

*\*RRI refers to the Relative Red Value; RGI refers to the Relative Green Value; RV, GV, and BV refers to the Red, Green, and Blue in the RGB values*.

The blue value is considered as a parameter because the intensity of the blue value will also affect the appearance of the color as camouflage in nature. The relative index is proportional, which means that the image takes from different strengths of light and positions of the nymph can be adjusted to the same level (Chang, *et al*., 1996).

## 3. Results

### 3.1 Hatching

The oothecae are in a shuttle shape, around 4 to 5cm. Its color is brown and full of alternate darker and brighter patterns. The entire hatching process passed through a month. The eyespots appeared in the 2^nd^ week, which indicates the oothecae is fertilized. Over 40 individuals are hatched from each ootheca. The very first ecdysis happened after hatch.

### 3.2 1st instar

The nymphs just hatched are in the 1st instar. They are about 3-4 mm long from the front to the end and 1 mm wide. The body color of the *Creobroter gemmata* at the time is unlike other stages. It consists of orange and black. Except for the pronotum, the latter half of its abdomen, the joint of the femur with the tibia of the grasping legs and the compound’s eye is black, and other parts of the body are orange. The food is *Drosophila melanogaster*; fed after the nymphs are just hatched. About ten flies are sufficient on a weekly basis.

During this period, a hypothesis about Batesium mimicry can be drawn out of the appearance of the 1st instar nymph. The mimic subject of the *Creobroter gemmata* nymphs in 1st instar probably is the *Tetraponera rufonigra*, a species of ants. The *Creobroter gemmata* 1st instar nymph shares several similarities with the worker of *Tetraponera rufonigra*:

1. The size of the nymph and the worker ant are both around 3-4mm.
2. The color of both nymph and worker ant is orange with black patterns.
3. *Tetraponera rufonigra* worker ants’ entire abdomen, femur, tibia, and head are black, which are corresponded to the black pattern of *Creobroter gemmata* nymph’s pronotum, abdomen, and the joint of the grasping leg.
4. *Creobroter gemmata* and *Tetraponera rufonigra* both distribute in the south and southeast Asia (Pfennig, *et al*., 2021; Nawphral, *et al*., 2021) (Fig. 3).

**Fig. 3.**
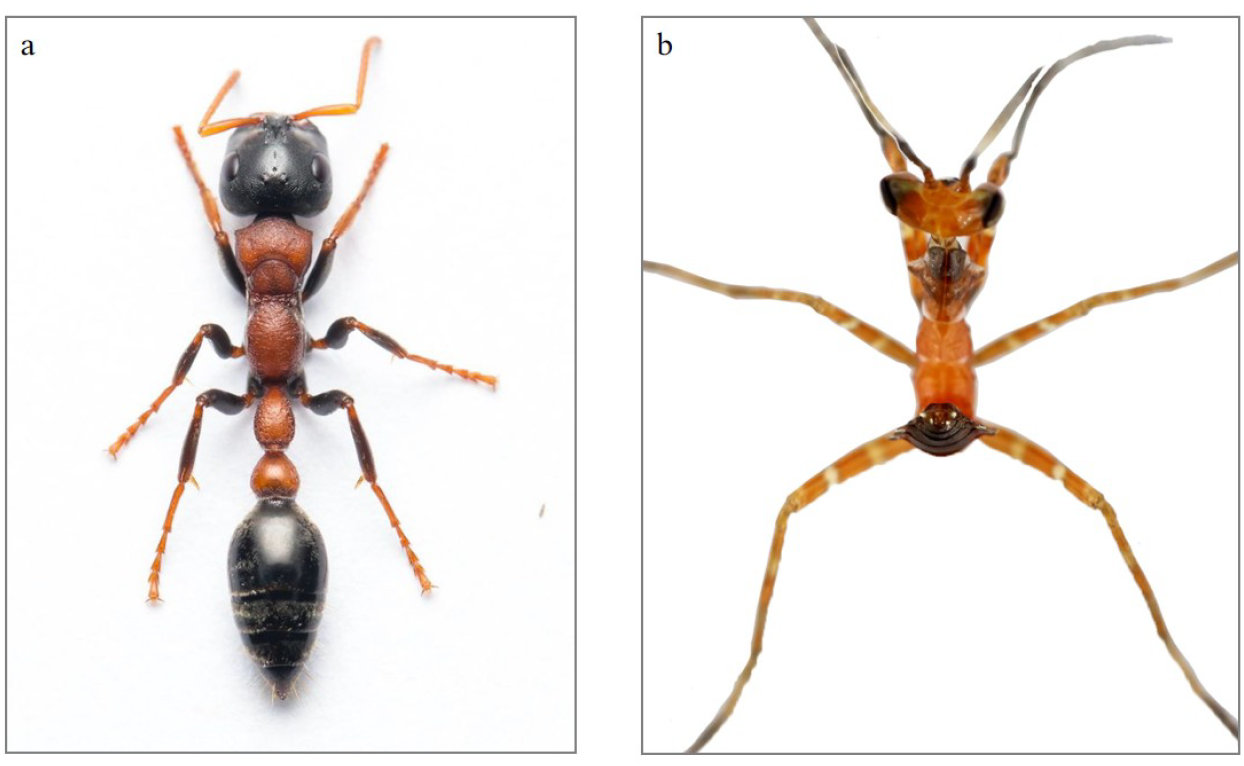
Picture (a) is the poisonous tropical worker ant *Tetraponera rufonigra*, and picture (b) is a *Creobroter gemmata* nymph in the 1st instar. This article suggest their similarity reflexes a type of Batesian mimicry.

Besides these, the strong toxicity of *Tetraponera rufonigra* also effectuates the protection by deterring predators.

### 3.3 2nd instar

After the first ecdysis, the 1st instar nymphs will become the 2nd instar nymphs. They are about 5-6 mm long and 2-3 mm wide. The pattern now is similar to the adult. Generally, the 2nd instar nymphs are slightly close to the 1st instar nymphs. Nymphs at 2nd instar need about 6-8 *Drosophila pseudoobscura* flies or over 20 *Drosophila melanogaster* flies per week (Fig. 4).

**Fig. 4.**
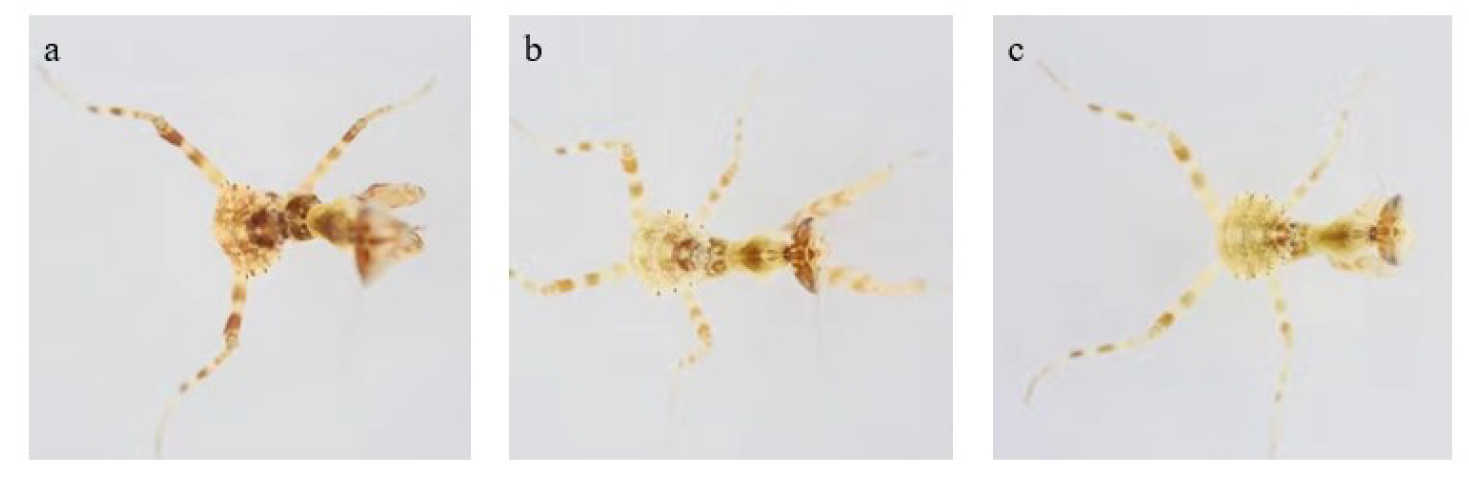
(a), (b), (c) are the pictures of *Creobroter gemmata* nymphs in 2nd instar, which were raised in (a) red box, (b) white box, and (c) green box.

### 3.4 3rd instar

After the second ecdysis, the 2nd instar nymphs will become the 3rd instar nymphs. They are about 8-10mm long and 3-4 mm wide. The body color is differentiated to conform with the background color. The difference of body color between individuals raised in the red and green background is the most evident in this stage (Fig. 5).

**Fig. 5.**
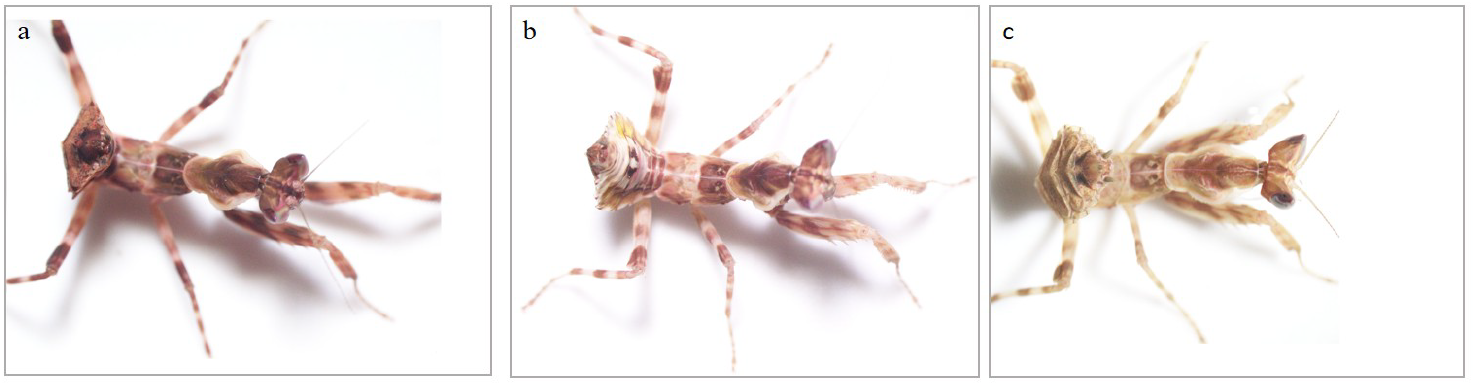
From the left to right are the pictures of Creobroter gemmata nymphs in 3nd instar, which were raised in (a) red box, (b) white box, and (c) green box. In this stage, the differentiation is the most significant.

Nymphs at the 3rd instar need about 10-15 *Drosophila pseudoobscura* flies per week.

### 3.5 4th instar

After the third ecdysis, the 3rd instar nymphs will become the 4th instar nymphs. They are about 10-15mm long and 5-8mm wide. Although the difference of body color between the red box group and the green box group is still visible, some individuals’ are slightly weakened. Nymphs at the 4th instar need about 18-20 *Drosophila pseudoobscura* flies per week.

### 3.6 5th instar

After the fourth ecdysis, the 4th instar nymphs will become the 5th instar nymphs. They are about 10-15mm long and 5-8mm wide. Individuals will present random body-color tendencies. Nymphs at the 5th instar need about 25-30 *Drosophila pseudoobscura* flies per week.

### 3.7 Photographing and data processing

Mantis pictures required an exposure rate from about 7.2 to 7.6. There should be sufficient light from the bottom and the top of the sample. The ideal body part of the nymph pictured is placed at the center of the graph. Since the purpose is to acquire the color of the nymph’s body parts, the other section of the entire graph is unnecessary to be as clear as its middle. The image quality of its center is ensured by feeding the nymph while taking photos. This method can effectively reduce its activity and therefore gain a picture of better quality (Kerr, *et al*., 2008). (Fig. 6)

**Fig. 6.**
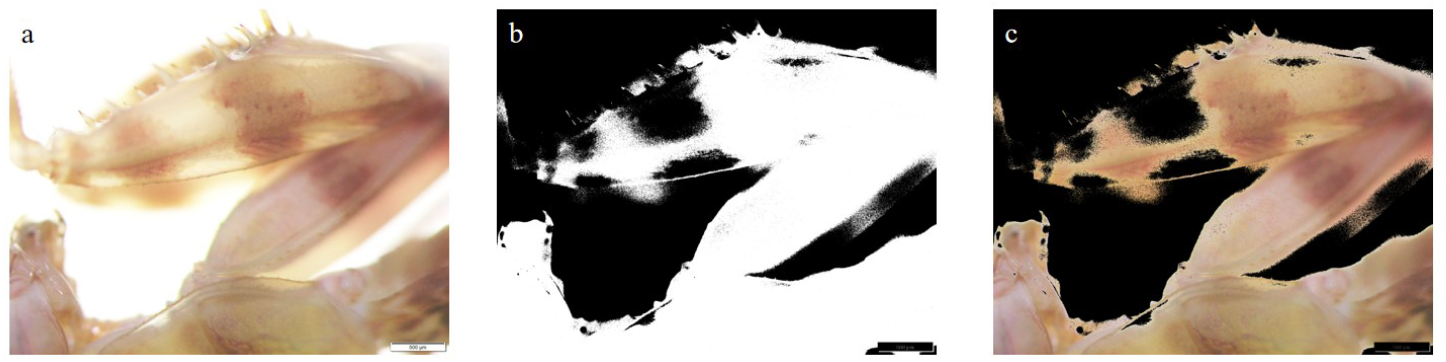
(a) is a picture of grasping leg of a 3^rd^ instar nymph from red box. (b) represents pixels the program selected through the thresh hold: only pixels with one value from RGB three values are lower than 230 or greater than 25 are valid. Black section are the pixels eliminated and white section are the pixels conserved. (c) is a picture constructed with eligible pixels.

Since the overly bright and overly dark color - cause by shade, over-exposure, or posture of nymphs - will dilute the significance of mantis body color, the value either too high or too low is not accepted. 230 and 25 are the thresholds for qualifying the pixels.

The value of color which is tested for the index is excluded in the denominator. This makes the result of color index into a directly proportional function, which can be better quantified and compared. Every picture will be fitted by taking the average of three values of RGD respectively into a single color block, which represents its general chromatic looks. The detailed data is recorded in a TXT file for further analysis (Figs. 7-8).

**Fig. 7.**
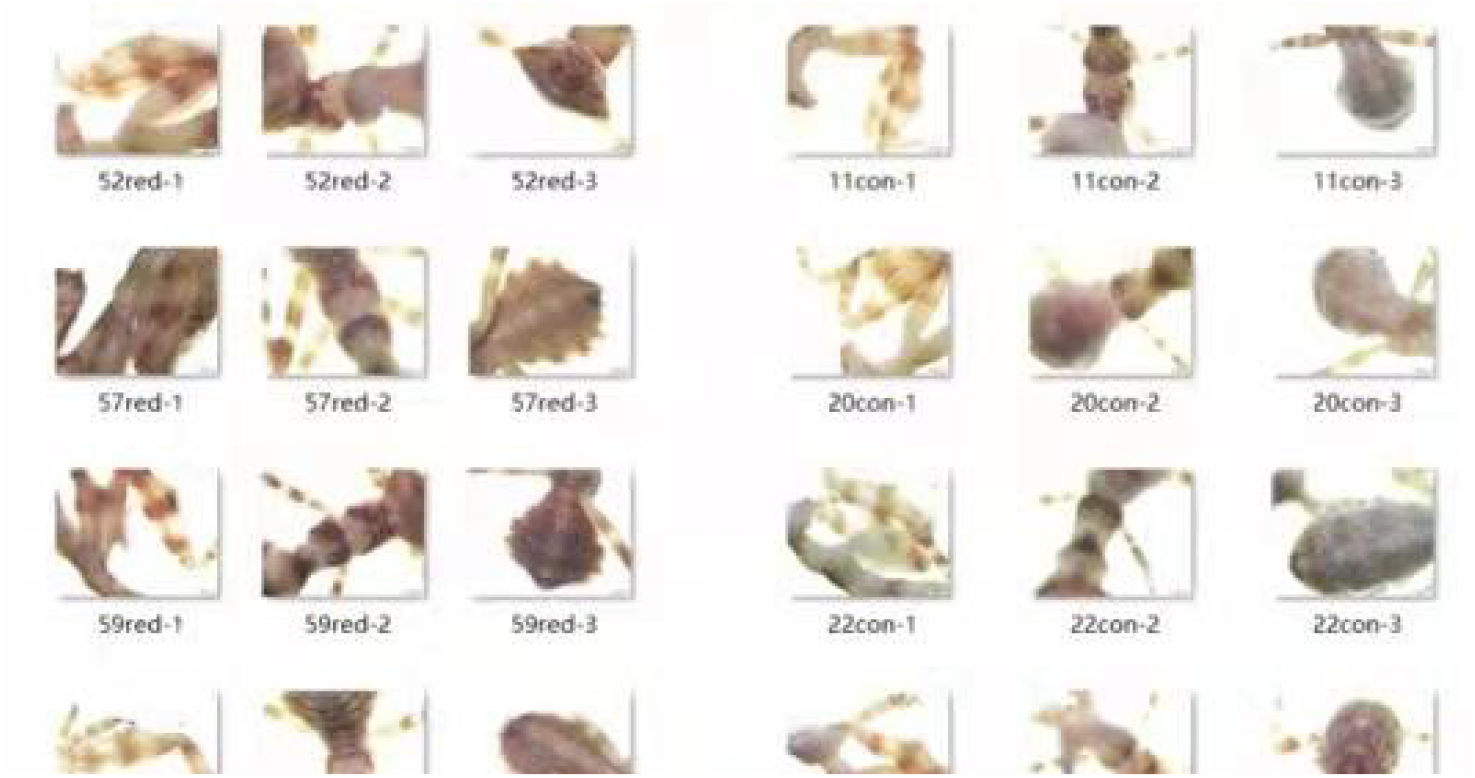
Pictures of the three significant body parts of the 5 individuals which were randomly selected from the 20 3rd instar nymphs of each group. Picture (a) is from the red box group, (b) from the control group, and (c) from the green box group.

**Fig. 8.**
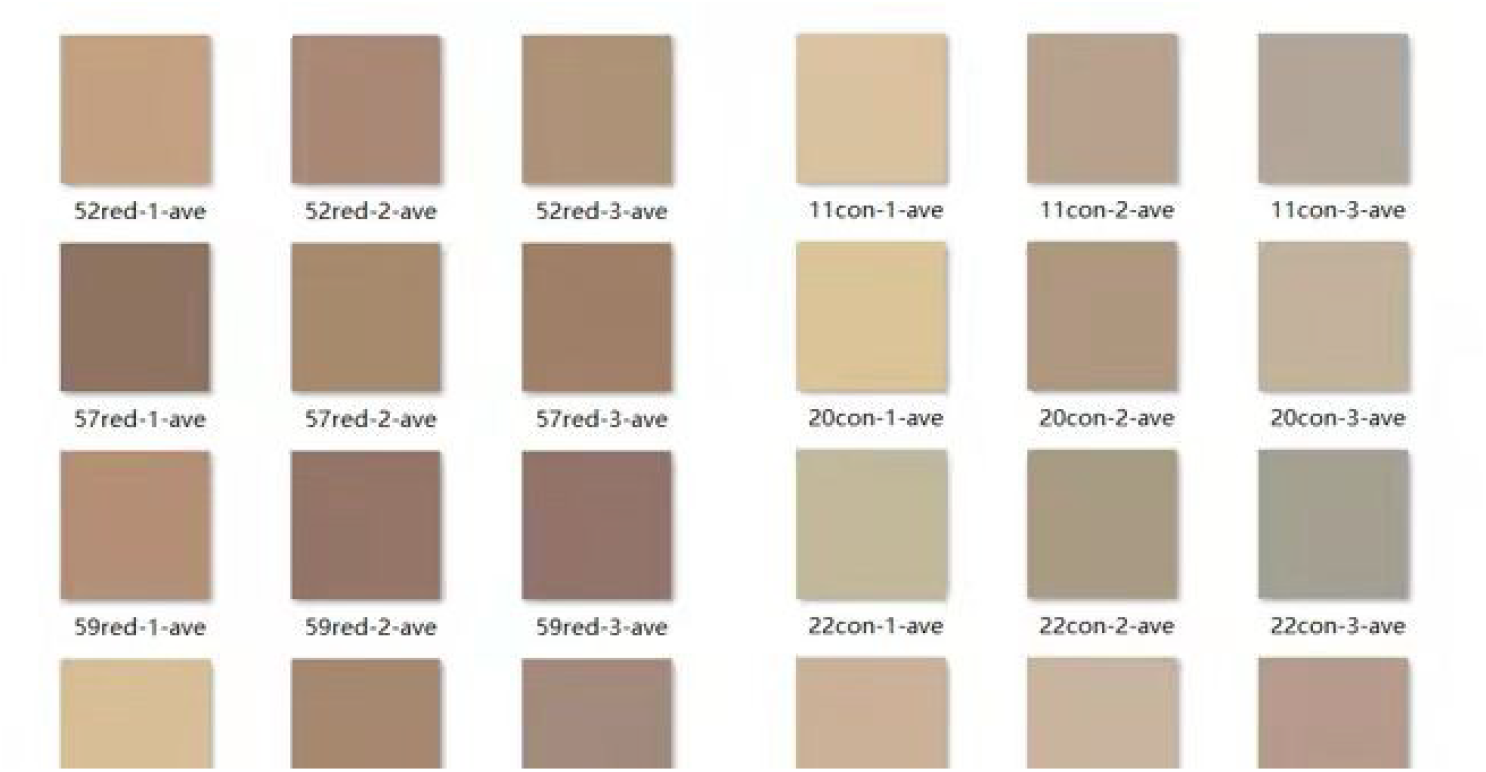
These are the color blocks fitted from the pictures in figure 10. Picture (a) is from the red box group, (b) from the control group, and (c) from the green box group.

### 3.8 Summarizing trend

The final data is taken from mantis nymphs from 1^st^ instar to the 5^th^ instar. The 1^st^ instar and 5^th^ instar contain twenty samples and sixty pictures for each group (the 1^st^ instar has only one group). The rest of the 2^nd^, 3^rd^, and the 4^th^ instar contain 5 samples and fifteen pictures in each group.

The data reveals that there is an effect on the body color of *Creobroter gemmata* from the background color. Generally, the differentiation between body color of nymphs raised in the red and green substrate is the most significant when they are at 3rd instar. The differentiation happens from the 2nd instar and gradually diminishes after the 3rd instar. The trend will be described separately at first, and then horizontally compared.

Start with the red background, the *relative red* of nymphs decreases at first from 67.52 to 64.20 after the first ecdysis, given that the initial color is lean to red, and then increases to the point of 67.33 after the second ecdysis. After the third ecdysis, the *relative red* decrease to 61.60 and stay still at 61.72 after the fourth ecdysis. It goes through a fluctuation at the 2nd instar and generally declines from the 3rd to 5th instar. Afterward, the relative green of nymphs decreases at first from 49.44 to 47.27 after the first ecdysis, and then it gradually increases to the point of 47.86 after the second ecdysis. After the third ecdysis, the relative green continually rises to 48.53 and stays still at 48.82 after the fourth ecdysis. So it goes through a declination at the 2nd instar and overall has a rise after then.

The second is the green background. The relative red of nymphs decreases at first from 67.52 to 63.20 after the first ecdysis, and then successively decreases to the point of 56.33 after the second ecdysis. After the third ecdysis, the *relative red* decrease to 52.73 then suddenly soar to 60.38 after the forth ecdysis. It goes through a fall from the 2nd to 4th instar and abruptly reverses its trend. Whereupon, the *relative green* of nymphs starts with an increase from 49.44 to 50.73 after the first ecdysis, and then it keeps on increases to the point of 51.06 after the second ecdysis. After the third ecdysis, the *relative green* continually rises to 51.33 and declines back to 49.46 after the fourth ecdysis. So it goes through a consecutive increment until it drops down at the 5th instar.

Lastly, as the control group with the white background, the *relative red* of nymphs decreases at first from 67.52 to 60.13 after the first ecdysis and then increases to the point of 60.87 after the second ecdysis. After the third ecdysis, the *relative red* decrease to 57.40 and stay still at 61.53 after the fourth ecdysis. So it keeps waving to a relatively small extent and generally declines from the 1st to 5th instar. Then, the *relative green* of nymphs decreases at first from 49.44 to 48.93 after the first ecdysis, and then it gradually increases to the point of 50.00 after the second ecdysis. After the third ecdysis, the *relative green* continually rises up to 50.80 and stays still at 48.82 after the fourth ecdysis with a small decline. So it goes through growth from the 2nd to the 4th instar but overall drops down (Fig. 9).

**Fig. 9.**
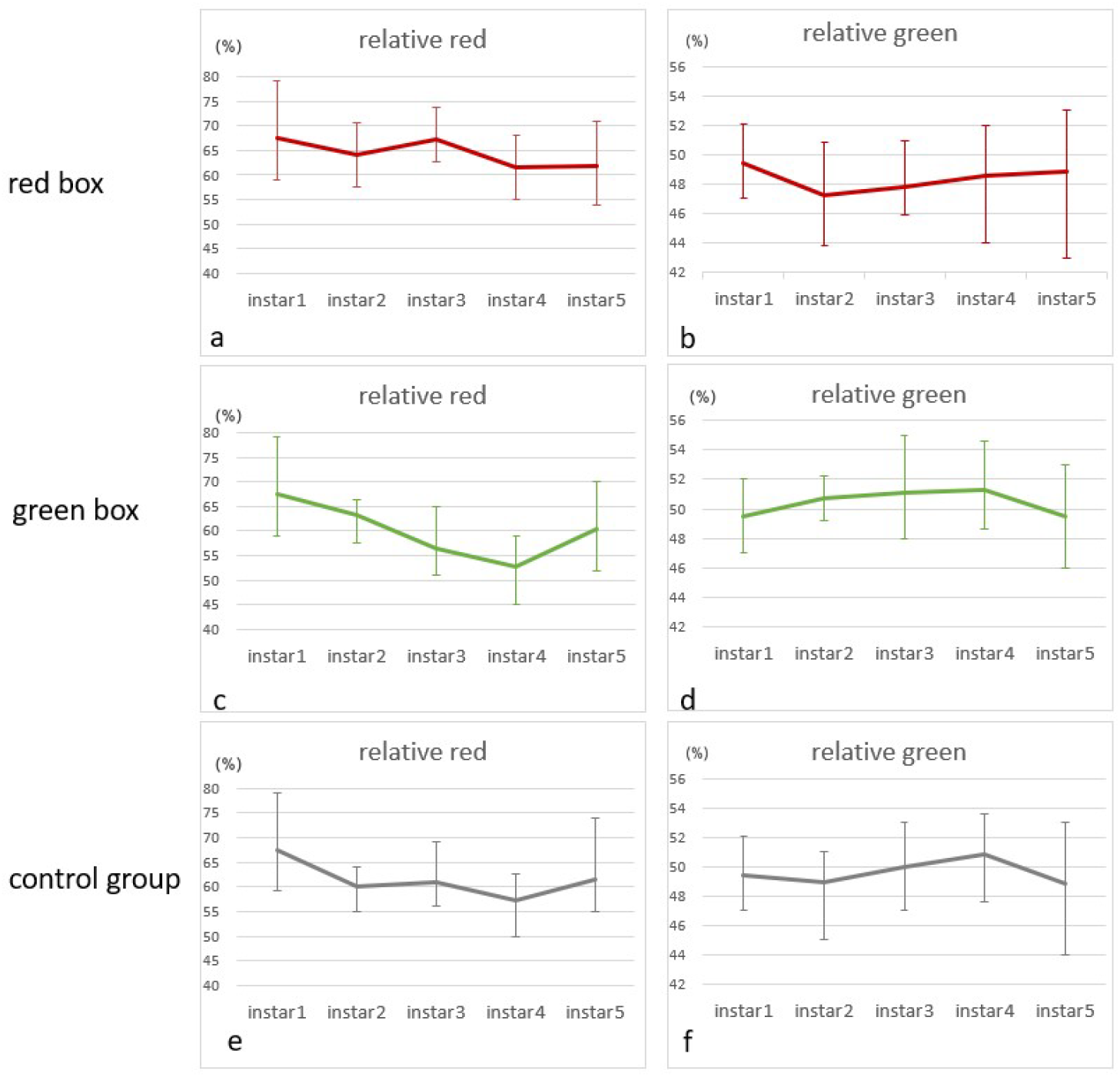
Three groups of *Creobroter gemmata* nymphs are separately raised in red box with red substrate, white box with no substrate, and green box with green substrate. White box is for control group. Graph (a),(c) and (e) show the *relative red* value of red box group, control group and green box group throughout the 1-5 instar. Graph (b),(d) and (f) show the *relative green* value of red box group, control group and green box group throughout the 1-5 instar. *\*Relatively red index = 100% * Red value / (Green value + Blue value)* ; *100% * Relatively green index = Green value / (Red value + Blue value)*.

The following is a comparison of indicators acquired from three different groups. It shows that the 3rd instar is the period when the differentiation is best revealed. (Test Relative Red: R-G 2.61108E-08, R-W 5.32173E-06, G-W 0.010131398; Relative Green: R-G 4.57114E-05, R-W 3.57714E-05, G-W 0.283044409). Compared horizontally, The differentiation of nymphs’ body color started from the 2nd instar, where the *relative green* has already shown distinction, and at the 3rd instar, both *relative red* and *relative green* get to a large extent of polarization. Thereafter, the difference starts to reduce and eventually about to vanish (Figs. 10-11).

**Fig. 10.**
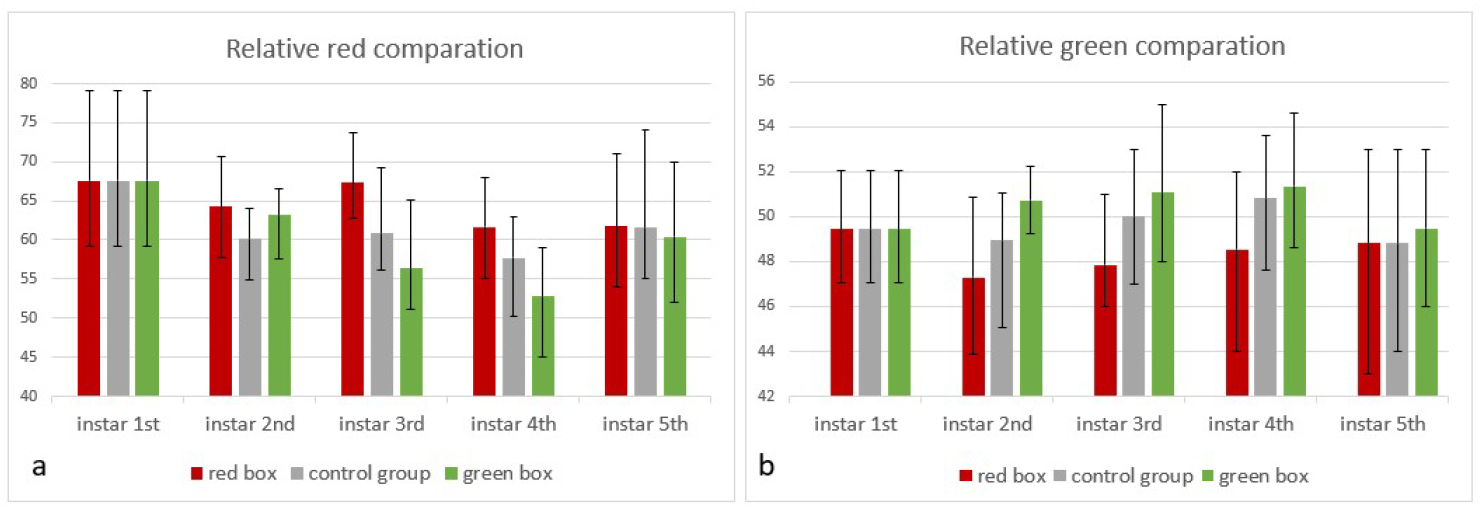
Three groups of *Creobroter gemmata* nymphs are separately raised in red box with red substrate, white box with no substrate, and green box with green substrate. White box is for control group. Graph (a) shows the histogram of *relative red* value of red box group, control group and green box group throughout the 1-5 instar for comparison. Graph (b) shows the *relative green* value of red box group, control group and green box group throughout the 1-5 instar for comparison. *\*Relatively red index = 100% * Red value / (Green value + Blue value)* ; *100% * Relatively green index = Green value / (Red value + Blue value)*.

**Fig. 11.**
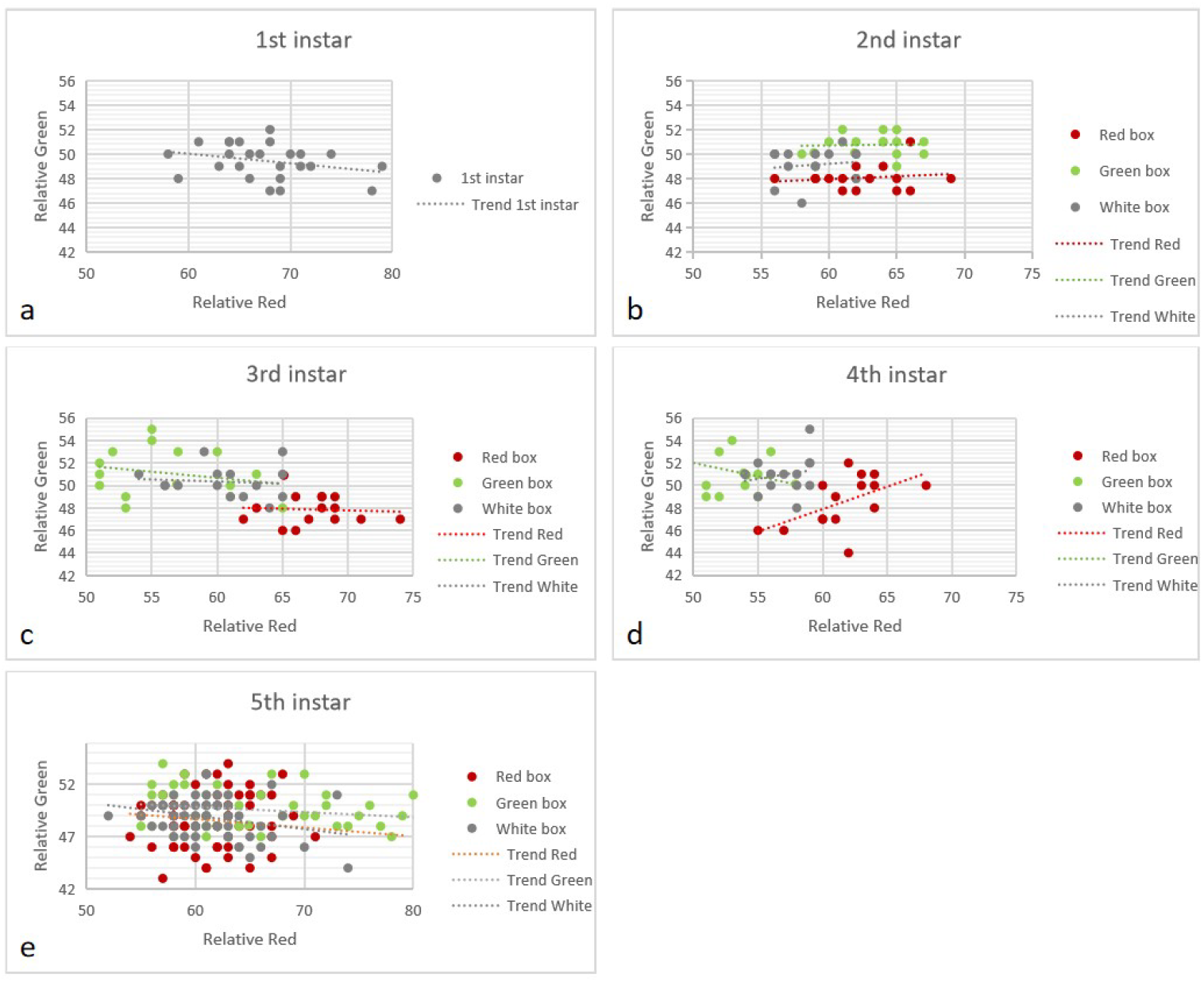
Three groups of *Creobroter gemmata* nymphs are separately raised in red box with red substrate, white box with no substrate, and green box with green substrate. Scatter graphs (a)-(e) show the distribution of *Relatively red* and *Relatively green* values in 1-5 instar. *\*Relatively red index = 100% * Red value / (Green value + Blue value)* ; *100% * Relatively green index = Green value / (Red value + Blue value)*.

**Fig. 12.**
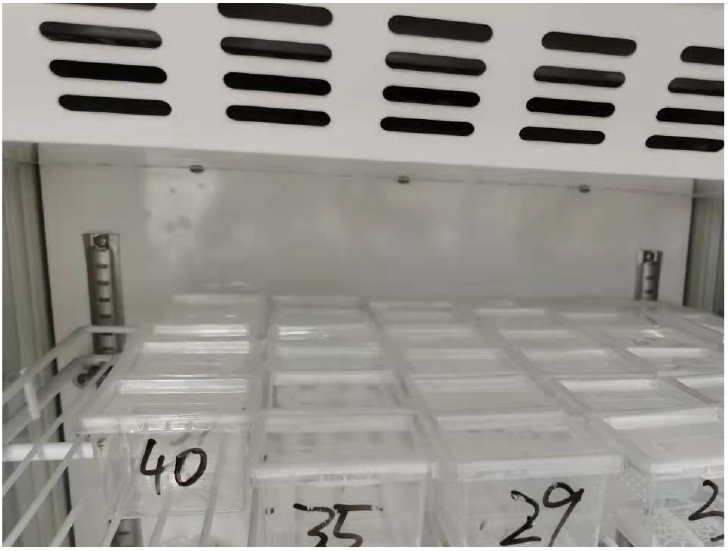
Incubator device

**Fig. 13.**
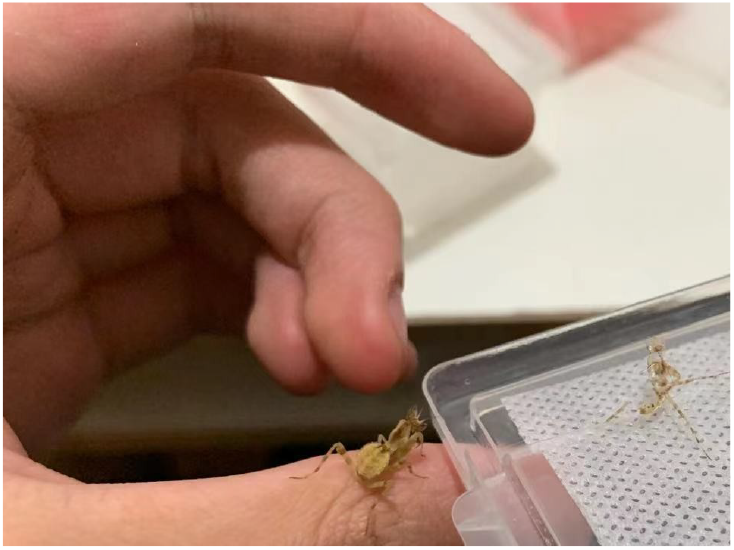
3^rd^ instar nymphs

**Fig. 14.**
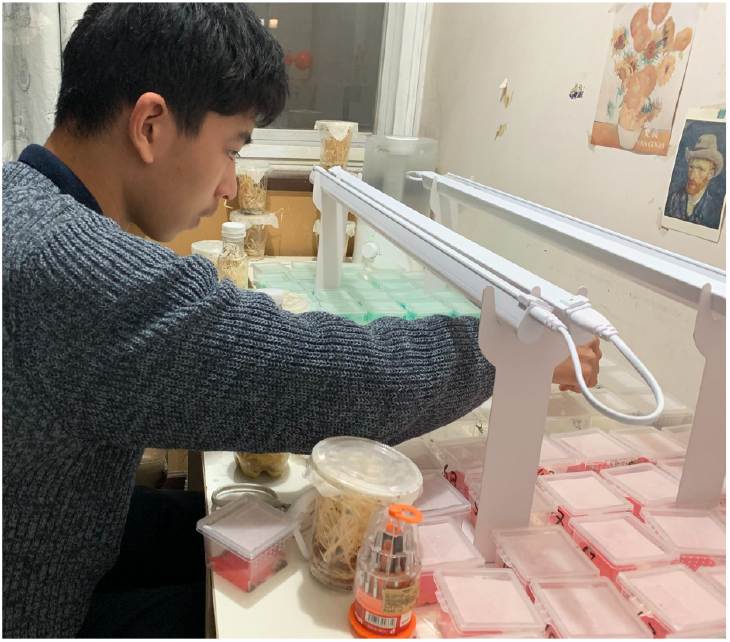
Winter holiday home-made lab with constant

**Fig. 15.**
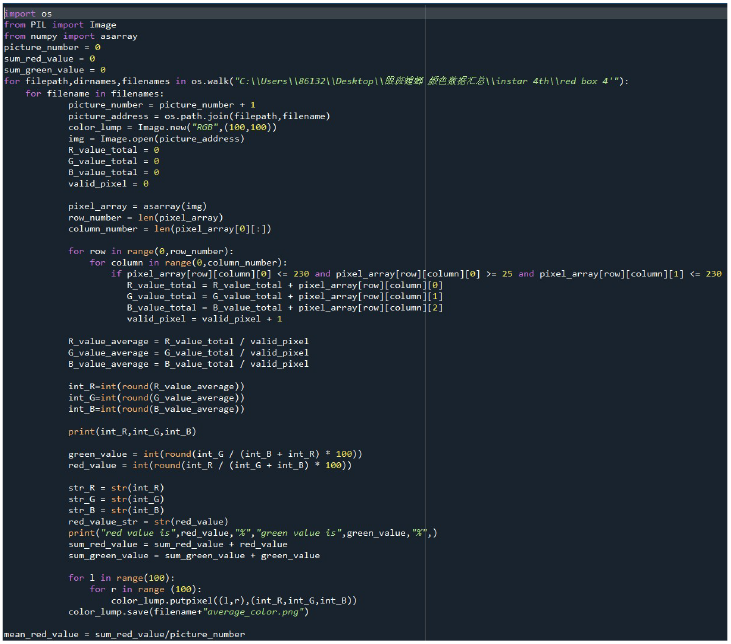
Program code written in python temperature, moisture, and optical period.

## 4. Discussion

This research uses image analysis and RGB values to quantify praying mantis’ camouflage in early stage, which is unprecedented. Previous researches about praying mantis mainly focused on their attractiveness as a flower towards the pollinating insects (O’hanlon, *et al*., 2013), their color pattern (Barabás, *et al*., 1999), and their wildland distribution. RGB values are also introduced as indicators to quantify insect body color for taxology by matching fixed sections (Lehnert, *et al*., 2011). Specifically, RGB values are suitable for this experiment, since red and green are two colors that are most present in *Creobroter gemmata’s* body color, and enable to compare the trend of insects body color in different instar in a linear relation. Formerly, RGB values are used to distinguish the difference between related species (Comeault, *et al*., 2016). The technique is maturely practiced by taxonomists and control tests (Joshi, *et al*., 2020). However, the vertical tracing of subject insects in different instar to acquire their body color differentiation trend is rarely revealed.

Previous researches are mostly focused on the horizontal comparison of subjects insects specific stages (Tanaka, *et al*., 2016). The background color also has been deemed as a variable for an experiment about insects’ crypsis (Hochkirch, *et al*., 2008), but the influence of background color on many species is barely documented. Substrate color is not used to conduct an experiment that is able to quantify its effect before. This experiment first time measured the influence of substrate color on mantis’ body color. Mimicry is one of the most common features of species under *Orthopterodea*, such as *Phasmatode*a, *Orthoptera*, and *Mantodea*. Among these species, *the Mantodea species* need mimicry most, because they not only use the camouflage to protect themselves as other insects do but use them as a tool to hunt. Throughout the evolution, the praying mantis which cannot perfectly hide in covert were eliminated by hunger. Moreover, the poor flying ability promoted the allopatric speciation of praying mantis and consequently formed myriad distinguishing specious.

Unlike the former belief that insects’ body color fully depends on their genetic material, and is passively selected by the environment, this experiment revealed another possibility of insects’ adaptation to the environment. *Creobroter gemmata* also behaves well as an experimental animal. Their small body size, short growth cycle, and multiform color pattern are all appropriate for the experiment. This research not only answered an intriguing question of whether some species of praying mantis will adapt their body color according to their surroundings but gives us some clue about the mechanism of acquired camouflage. Inconsistent with the general knowledge that the body color of a mantis is determined by the genetic for the most part, nurture also has played a big role in mantis morphology (Wittkopp, *et al*., 2009). This point has been proved in many experiments of grasshoppers but never applied on mantis species (Edelaar, *et al*., 2017). The substrate color of red is also uncommon in previous researches in insects because the grasshoppers usually are green or brown. Nonetheless, flower mantises live on the stem or foliage of plants, so they possess brighter and more colorful body colors, like red, purple, or orange (Koh, 2020).

The trend of differentiation of nymphs’ body color in different instar is seldom recorded, and the result is interesting. The value of the indexes of *relative red* and *relative green* of *Creobroter gemmata* showed a preference for the red color, unlike most insects life in leaves, which is corresponded to their flowery, habitat. And the trend also reveals a camouflage adaptation from stage to stage. The inclination of body color to match circumstances begins and is significant in the early stages of the nymphs: the 2nd and 3rd instar, but start to unify in the latter stages. It comes up with a hypothesis: The early stages of nymphs are too small and weak to protect themselves from the attack from predators so they need some techniques to prevent the potential danger. As soon as the nymph hatched, they have no time to adjust their body color to adapt to surrounding colors, so that they mimic a toxic ant species *Tetraponera rufonigra* to scare off predators. After ecdysis once, nymphs become too big to mimic ants, so they will hide in the foliage and flowers or mimic as a part of plants with their adapted body color. For example, if the nymphs live in a cluster of bushes or leaves, it will tend to be green, or if it lives in a plant with plenty of pinky flowers, it will tend to be red. The early stages need more cautious camouflage to enhance surviving rates. However, after the 4th instar, the nymphs will become nearly 15mm long, which is big enough to tackle with most of the predators in its habitat (Fontanilla, *et al*., 2019), and spread out from the place it originally hatched and stay. In this case, the habitat might become different from where they adapted to, so a medium color that is suitable for most of the surroundings is fixated on their body color. And this is how the adjustment of their body color in the early stages may work.

Sample quantity has been one of the biggest problems of the research. Mantis as an experimental subject is uncommon, and unlike other herbivore insects, rearing mantis requires separated enclosures and feeding, which will cost much more time and space. Those factors make it hard to provide as many samples as other samples from model organisms, like the grasshopper (Tanaka, *et al*., 2012). In this experiment, it was expected to have 30 samples for each comparative group at first. However, the limitation of space and high rate of died young result in only 20 individuals were survived and recorded. Except for the first and the fifth instar, others have only 5 randomly selected samples. The short time interval between the early instars and the insufficient experimental hours results in a limitation in the number of graphical supply. On the other hand, the laboratory in the school was unavailable during the winter and summer vacation due to the COVID-19 pandemic also made it impossible to repeat the mantis’ nymphs keeping process and collect extra data. During the winter vacation Nymphs had to be kept in a homemade condition. Although the environment had gone through a slight change from the incubator in the school lab to the homemade site, the moisture, temperature, light period, and light intensity were as much as possible to be consistent with the former environment. Although the experiment might become less convincing due to the lack of sample size, the qualitative results are still valuable.

The experiment is a super-facial exploration of the field of mantis’ camouflage (Pembury, *et al*., 2020). There are still many delving-worthy perspectives of the research. From the biochemical point of view, insects possess various types of pigments, including anthraquinones, aphins, pterins, tetrapyrroles, ommochromes, melanins, and papiliochromes. Tetrapyrroles consist of four pyrrole rings, connected by one-carbon (methine or methylene) bridges, in either a linear or cyclic manner. Bilirubin and phycobilin are linear tetrapyrroles (bilanes) with three one-carbon bridges and porphyrins and chlorins are cyclic tetrapyrroles with four one-carbon bridges. This pigment was found in Mantodea as a green pigment (Shanim, *et al*., 2014). Red pigments also played a significant role in the mantis’ background color adaption. The major color components of insects’ pigments are carminic acid, kermesic acid, and laccaic acids, respectively. These dyes are red anthraquinone derivatives (Cooksey, 2019). Although red pigment is uncommon to see on species in Orthopteroidae, *Creobroter gemmata* large probably can produce one of the above red pigment molecules. Behind the phenomenon of body color adaptation, a complete colored-light feedback pigment synthesis system should exist and be unknown (Briscoe, *et al*., 2001). In the further molecular biology experiment and investigation, a background color responding system including sensor, transmitting material, and pigment generator should be unveiled. In this hypothesis, *Creobroter gemmata* should possess certain apparatus to receive different wavelengths of light. Praying mantis’ compound eyes are eligible organs for detecting colors in the spectrum (Sontag & Charles,1971). Pigment-dispersing hormones are proved to be a transmitting factor to control insects’ body color during their life cycle (Helfrich-Forster, 2005). The final receptor of the entire system is a pigment producer. Pigments are produced by epidermal cells through a developmental process that includes pigment patterning and synthesis (Wittkopp, *et al*., 2009). As the compound eyes of *Creobroter gemmata* receive light in a certain wavelength of green or red, it may stimulate the endocrine glance to secrete pigment-dispersing hormone. Then, epidermal cells will synthesis of red or green pigment according to the hormone. This process should be more significant when the stage of the 1st instar and the 2nd.

The phenotype of background color adaptation present in *Creobroter gemmata* should relate to a certain genotype. These genes can help Praying mantis hide better in the early stages and increase the surviving rate of individuals (Fukutomi, *et al*., 2021). Future research can focus on whether the ability of substrate color imitating is hereditary and which genetic locus controls it.

## Acknowledgement

Great thanks to Dr. Siqin Ge from the Institute of Zoology, Chinese Academy of Sciences, and Dr. Feng Li from RDFZ for their voluntary support and advice on experiment design, facilities, and data analysis.

The idea of the research is come from Shao Zhi Geroge Liu’s praying mantis keeping experience and internet forums. The experiment was personally initiated and executed, and the paper was written with advice and polish from Dr. Siqin Ge.

Experiment design: Shao Zhi George Liu

Experiment operation: Shao Zhi George Liu, Renzhi Ma

Data analysis: Shao Zhi George Liu

Results and conclusion: Shao Zhi George Liu

Paper writing: Shao Zhi George Liu

## Appendix

